# Long-term stomatal and leaf trait dynamics in invasive knotweeds in Europe - insights from 160 years of herbarium records

**DOI:** 10.64898/2026.07.08.737080

**Authors:** Felix A. Hahn, Franziska M. Willems, Luzia Hamma, Farah Badreldin, Christiane Karasch-Wittmann, Ann Bogaerts, Madalin Parepa, Uta Grünert, Christina L. Richards, Oliver Bossdorf, Ramona-Elena Irimia

## Abstract

1. Stomata and leaf traits are key regulators of plant water use efficiency and are expected to have changed in response to rising atmospheric CO_2_ concentrations and climate warming over the past centuries. However, long-term data documenting such changes are rare.
2. We leveraged herbarium collections to track changes in stomatal characteristics and leaf traits in 656 individuals of invasive Japanese knotweed and its hybrid Bohemian knotweed collected across their European range and spanning 160 years of invasive spread.
3. We found that several functional traits including stomatal density and maximum anatomical stomatal conductance did not show significant changes over time but that plants adjusted their stomatal size and shape over time, and that these changes were associated with increased atmospheric CO_2_ levels. Interestingly, *Reynoutria japonica* showed increases in stomatal size and stomatal elongation, while the hybrid *R.* × *bohemica* showed a reduction in stomatal size. Traits also varied systematically with climates of origin. Plants from warmer origins with higher evaporative demands during the growing season had thicker leaves, lower SLA, smaller stomata and higher stomatal density, indicating more conservative water-use strategies. Stomatal density and gas exchange capacity co-varied with leaf structural traits, and there was a trade-off between stomatal size and number. Overall, “fast” leaf economic traits were associated with “slow” physiological traits.
4. Our results suggest that stomatal anatomical plasticity may enhance climate resilience by maintaining a stable maximum gas exchange capacity across environmental gradients. Herbarium collections provide a unique resource for reconstructing plant responses to historical environmental changes and understanding intraspecific trait variation.

## Introduction

Atmospheric CO_2_ has risen from ∼280 ppm in the pre-industrial era to ∼420 ppm currently, significantly altering terrestrial ecosystem conditions. Elevated CO_2_ often increases photosynthesis and water-use efficiency and can favor fast-growing, resource-acquisitive species. This can shift plant community composition by changing species dominance, reducing biodiversity in some systems and promoting the expansion of groups such as woody plants and invasive species (Ainsworth and Long 2005; Mndela et al., 2022; Sobuj et al., 2024; Cadotte et al., 2026).

A variety of morphological and physiological traits contribute to plants’ response to environmental change. The framework of the leaf economic spectrum describes a trade-off between resource acquisition and conservation: species with “fast” strategies invest in rapid growth and high resource turnover (e.g., high specific leaf area, SLA), while “slow” strategies prioritize leaf longevity and resource conservation (Wright et al., 2004). Within this framework, leaf size influences boundary-layer thickness and thus thermal and water regulation, while stomata control CO_2_ uptake and transpiration and thus jointly shape overall plant physiological performance. In addition, variation in stomatal density determines pore number per unit leaf area and therefore potential maximum conductance, whereas stomatal size influences maximum conductance and the speed of stomatal response, with consequences for photosynthesis, water-use efficiency and ecosystem productivity (Franks and Beerling 2009; Hoffman et al., 2025). In many cases, fast economic traits (e.g., high SLA) are associated with fast hydraulic traits—such as small, densely packed stomata—supporting high photosynthetic rates and rapid water transport (Yin et al., 2018). However, this coordination is not universal. Recent studies show that leaf economic strategies and hydraulic traits can be decoupled: for example, some species exhibit fast growth strategies alongside slow hydraulic traits, such as fewer, larger stomata that limit water loss but may constrain photosynthetic capacity (Pan et al., 2024; Liu et al., 2020). This suggests that trait coordination is context-dependent and shaped by environmental constraints.

Stomata are sensitive to various environmental change factors such as ambient CO2, temperature, light intensity, nitrogen deposition, reduced precipitation and ozone pollution. Consequently, studying their long-term dynamics offers valuable insights into how ecosystems adapt and respond to a changing climate (Liu et al., 2022; Liang et al., 2023). Stomata can respond to the environment on both short and long time scales, from minutes to millennia, through rapid opening and closing as well as gradual changes in their size, density and geometry (Haworth et al., 2011). It is also commonly observed that smaller stomata are often accompanied by higher stomatal density, indicating a compensatory relationship between size and number (Hetherington and Woodward, 2003). Across diverse biomes and plant functional groups, the most consistent response to elevated atmospheric CO_2_ appears to be a reduction in stomatal conductance (Liang et al., 2023). However, more plastic traits, such as stomatal density can exhibit complex and context-dependent responses, reflecting a trade-off between maximizing carbon uptake and minimizing water loss under varying environmental conditions (Yan et al., 2017).

Stomatal architecture can vary across climatic gradients. For instance, populations of *Andropogon gerardi* originating from arid regions developed smaller, denser stomata, a trait likely facilitating more rapid and dynamic stomatal regulation (Sytsma et al., 2025). Conversely, populations of *Arabidopsis thaliana* from cooler or wetter environments generally displayed fewer, larger stomata accompanied by reduced stomatal conductance (Elfarargi et al., 2023). Stomatal distribution across leaf surfaces further modulates these strategies: amphistomy (stomata found on both leaf surfaces) enhances conductance and carbon gain in high-light environments, while hypostomy (stomata primarily found on the lower leaf surface) reduces water loss in shaded, cooler or drier conditions (Drake et al., 2019). Understanding long-term changes in stomatal traits in natural populations can help inform predictions about plant and ecosystem responses to climate change (de Boer et al., 2011; Liang et al., 2023). Herbarium collections provide an effective resource to track shifts in phenotypic traits, phenology and species distribution across time and space (Heberling, 2022). Despite temporal and spatial gaps, these specimens uniquely enable the study of species-level responses over decades or centuries, helping to clarify patterns that short-term studies may miss.

*Reynoutria japonica* and its hybrid *R. × bohemica* are invasive herbaceous perennials widespread across central and western Europe, commonly found in riparian zones, disturbed sites and along roads, typically in open, high-light environments. Their dense growth and high leaf area can significantly influence the hydrologic cycle by increasing water uptake and transpiration (Cummins 2013; Galster and Vanderklein 2023; Adler-Colvin et al., 2025). Shifts in physiological function and phenotypic plasticity under varying environmental conditions may help explain their rapid and widespread expansion (Richards et al., 2008; Zhang et al., 2016; Wang et al., 2025; Irimia et al., 2026a). Here, we used a herbarium-based approach to examine the response to rising C02 and climate factors in introduced *Reynoutria* sp. over the past 160 years. We expected stomatal size to respond to rising CO2, potentially in coordination with stomatal density, such that maximum anatomical gas exchange capacity (g_max_) remains relatively buffered over time. To disentangle the drivers that could underlie these patterns, we then tested the effects of atmospheric CO_2_ and, separately, a suite of climatic variables to better understand trait co-variation. In addition, we tested whether the hybrid *R.* × *bohemica* differs from *R. japonica* in mean trait values and in its sensitivity to environmental conditions. We hypothesized that the hybrid would exhibit greater phenotypic diversity because of its higher genetic diversity (Tiébré et al., 2007; Krebs et al., 2009; Jugieau et al., 2024), broader environmental responsiveness, and higher mean physiological capacity (e.g., g_max_ and SLA). Finally, we examined the coordination between stomatal and leaf economic traits, to test for trade-offs reflecting how plants allocate resources between structural investment and gas exchange capacity.

## Materials and Methods

### Study species

Invasive knotweeds (*Reynoutria*, Polygonaceae) form a species complex comprising some of the world’s worst invasive plants (Lowe et al., 2004). The genus includes *R*. *japonica* Houtt. (Japanese knotweed), *R*. *sachalinensis* Nakai (Giant knotweed), hybrids between the two (*R.* × *bohemica* Chrtek & Chrtková, Bohemian knotweed) and crosses and backcrosses among the various taxa. Native to SE Asia, the parental species were introduced to Europe and North America from Japan in the 19^th^ century (Bailey and Conolly 2000). The taxa spreads predominantly through rhizome pieces or stems, but can also produce seeds when pollen becomes available (Bailey et al., 2009; D’Hertefeldt et al., 2026). *Reynoutria japonica* and the hybrid *R.* × *bohemica* are widely distributed across the Northern hemisphere and highly invasive, whereas *R. sachalinensis* is less common and has a reduced ecological impact (Bailey et al., 2007). Across Western Europe and the British Isles, populations of *R. japonica* are widely considered to comprise clonal replicates derived from a single octoploid female genotype (Zhang et al., 2004; Irimia et al., 2025; 2026b), whereas *R*. × *bohemica* hybrids are much more diverse and comprise both sexes and unique genotypes (Tiébré et al., 2007; Krebs et al., 2009; Jugieau et al., 2024).

### Herbarium phenotyping

We borrowed herbarium specimens of invasive knotweeds from 30 herbaria across Europe, to our herbarium in Tübingen, Germany. We determined the taxonomic identity of the specimens using the taxonomic keys of Zika and Jacobson (2003), and stored the specimens in fireproof metal cabinets at room temperature until phenotyping. Only specimens with recorded collection dates (at least month and year) and collection localities were included in this study. Additionally, we recorded each specimen’s barcode, collector name and habitat. We classified sites into major habitat groups reflecting both environmental setting and disturbance regime such as: coastal, riparian, urban managed, rural managed, natural or semi-natural, and disturbed/ruderal habitats (Fig. S1). We inferred elevation (m) for each specimen from latitude and longitude using a digital elevation model (DEM; Schneider 2019). We excluded specimens that were glued entirely to the sheet, as well as those with folded leaves that could be damaged during sampling. We phenotyped 656 specimens collected across Europe (Fig. 1), spanning climatic conditions from warm to semi-arid and sub-humid regions (Fig. S2). The dataset included 454 *R. japonica* and 202 *R. × bohemica*, sampled between 1861 and 2021. Of these, 39 specimens were collected prior to 1900, 99 between 1900 and 1940, 234 between 1940 and 1980, and 284 between 1980 and 2021.

**Figure 1.**
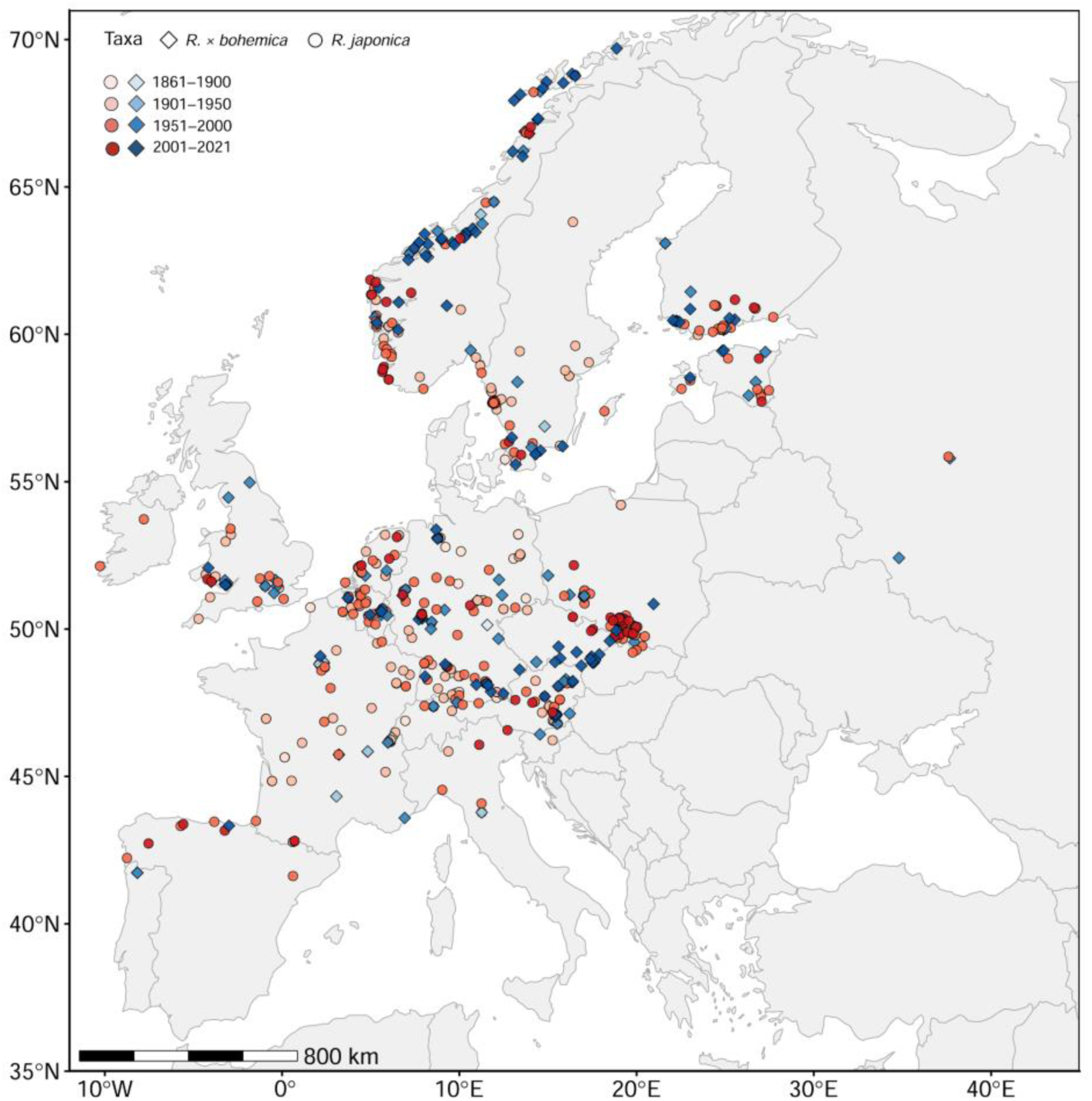
Geographic origins of the 656 samples (454 *R. japonica* and 202 *R.* × *bohemica*) included in this study. Color gradients indicate different sampling periods.

For each specimen, we measured specific leaf area (SLA), stomatal density and stomatal size. We attempted to include *Reynoutria sachalinensis* in the sampling, however, the stomata were too small and poorly visible to be reliably counted at the same magnification we used for the other taxa. In addition, the leaves of *R. sachalinensis* were very thin and brittle, and the stomata impression process frequently caused damage to the specimens. Consequently, we excluded *R. sachalinensis* from the study. To measure SLA, we selected two fully expanded mature leaves per specimen (the fourth and fifth or fifth and sixth leaf counted from the top) and detached one leaf (including the petiole) left of the stem and one leaf right of the stem using a sharp razor blade. We mounted the leaves onto a blank A3 paper and took pictures using a Sony RX 100 digital camera, then weighed each leaf to the nearest mg using a Sartorius Quintix 224-1S laboratory high precision scale. We used the average area and average weight of the two leaves per specimen to calculate the specific leaf area (mm^2^mg^-1^) and leaf mass per area (LMA: leaf dry mass/leaf area, mg^-1^mm^−2^). To determine the area of each leaf, we used ImageJ version v1.53.u.

We used a dental resin technique (AFFINIS^TM^, 50 ml, Coltene) to obtain stomatal impressions (Dunn et al., 2019). AFFINS is a two-component, addition-curing polyvinylsiloxane material commonly used for dental impressions. The base and catalyst were supplied in a cartridge and mixed in the correct ratio using a universal mixing tip, then applied with a dispensing gun. We applied the resin paste to the abaxial (lower) leaf surface of the two detached leaves, using a piece of plastic film with a punched hole. To avoid major veins, we placed two films on each leaf - one on either side of the central vein - approximately at the center of each leaf half in areas with minimal venation. Preliminary tests on one-year old dried *Reynoutria* leaves showed no significant differences in stomatal density among the upper, middle and lower parts of the leaf (Badreldin, unpublished). These tests also confirmed that *Reynoutria* sp. are hypostomatous, meaning that they bear stomata mostly on the abaxial leaf surface. During sampling, we applied an approximately pea-sized amount of resin to the hole in the plastic film and gently pressed it onto the leaf using a plastic tag to ensure proper contact. After approximately 15 minutes, the resin had set and we carefully removed the plastic film. We peeled the resulting mold off and stored it in a labeled paper bag. We repeated this process to obtain four molds per specimen. After these measurements, we re-attached the sampled leaves, undamaged back to the specimen.

To create epidermal imprints, we applied clear nail polish to each resin mold. After approximately 20 minutes, we peeled the dried nail polish layer off with tweezers and mounted the peel on a microscope slide with a coverslip secured using small drops of nail polish. We captured microscopic images using a Canon EOS 650D camera mounted on a Carl Zeiss Axioskop 2 Plus microscope using Canon EOS Utility 2 software with a 20× objective. We took one image per imprint, resulting in four images per specimen.

We determined stomatal density (number of stomata per mm^2^) using ImageJ. We marked stomata manually and they were automatically counted using the “Cell Counter” plugin. All observers followed a standardized counting protocol and included only clearly identifiable stomata. Because each image covered an area larger than 1 mm^2^, we divided stomatal counts by 1.12 to obtain density per 1 mm^2^, and calculated the mean stomatal density across the four images per specimen to account for within-individual variation. We measured stomatal complex size (including guard cells and stomatal pore area) from images captured at 40× magnification, on the same epidermal imprints as for stomatal density. We measured length and width of the stomatal complex (in µm) in ImageJ using a stage micrometer for scale calibration. For each image, we selected two stomata with clearly visible guard cell boundaries, resulting in a total of eight measurements per specimen.

### Statistical analyses

We obtained climate data from the Climatic Research Unit dataset (CRU TS4.06), extracting the following monthly variables for the year of collection of each specimen: average temperature (tmp, °C), average monthly maximum temperature (tmx, °C), total precipitation (pre, mm month^−1^), vapour pressure (vap, hPa), and potential evapotranspiration (pet, mm month^−1^). To better reflect biologically relevant conditions, we restricted climate data to the growing season, which was defined regionally as follows: northern Europe (Denmark, Norway, Finland, Estonia, Sweden, Ireland and the United Kingdom: May–September), central Europe (Austria, Belgium, Czech Republic, Germany, Netherlands, Poland, Slovakia, Slovenia and Switzerland) and southern Europe (France, Spain, Portugal and Italy) from April to September. For each specimen, we calculated growing-season means for temperature (tmp, tmx), vapour pressure (vap) and potential evapotranspiration (PET) and summed precipitation. For specimens collected prior to 1901, when CRU climate data are unavailable, we used mean values from 1901–1905 as a proxy for growing-season conditions across all variables. We also obtained atmospheric CO_2_ concentrations (ppm) from in situ measurements at the Mauna Loa Observatory for specimens collected between 1958 and 2021 (Keeling et al., 2005). For specimens collected before 1958, CO_2_ data were taken from ice core reconstructions provided by CSIRO (Rubino et al., 2019).

We calculated a series of standard composite stomatal metrics (Table 1). Stomatal complex area was calculated as:

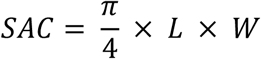

where *L* is guard cell length and *W* is the combined width of the guard cells and stomatal pore. Stomatal aspect ratio was calculated as:

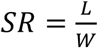

which describes the elongation of the stomatal complex (e.g. when length > width, stomata are long and narrow, when length ≈ width, stomata are more rounded or circular). Stomatal area fraction (*f*) was calculated as:

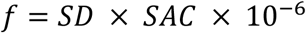

where SD is stomatal density (mm^−2^) and SAC is stomatal complex area (µm^2^). This metric represents the fraction of the leaf epidermis occupied by stomata. The theoretical maximum anatomical stomatal conductance to water vapour (*g*_max_) was calculated following equation by Franks and Beerling (2009):

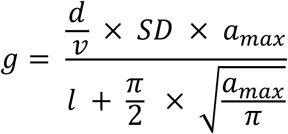

where *d* is the diffusivity of water vapour in air at 25°C (0.0000249 m^2^ s^-1^), *v* is the molar volume of air (0.0245 m^3^ mol^-1^), *l* is the stomatal pore depth (assumed to be equal to half of the guard cell length) and *a*_max_ is the maximum pore area. We calculated maximum pore area (*a*_max_), as the product of pore length and pore width. We approximated pore length as half the guard cell length, and pore width as half the stomatal complex width. We converted both measurements from micrometers to ensure unit consistency, giving *a*_max_ in square meters (m²) to calculate maximum anatomical stomatal conductance (g_max_). g_max_ is a structural estimate of the theoretical maximum gas exchange capacity, based on stomatal size, density and geometry, assuming fully open stomata under ideal conditions (Franks et al., 2009). Therefore, g_max_ is typically higher than physiological stomatal conductance (g_a_), which reflects actual, environmentally regulated gas exchange. We use g_max_ to define structural limits and physiological constraints on plant gas exchange capacity (Franks and Beerling 2009; Franks et al., 2009; de Boer et al., 2011).

**Table 1.**
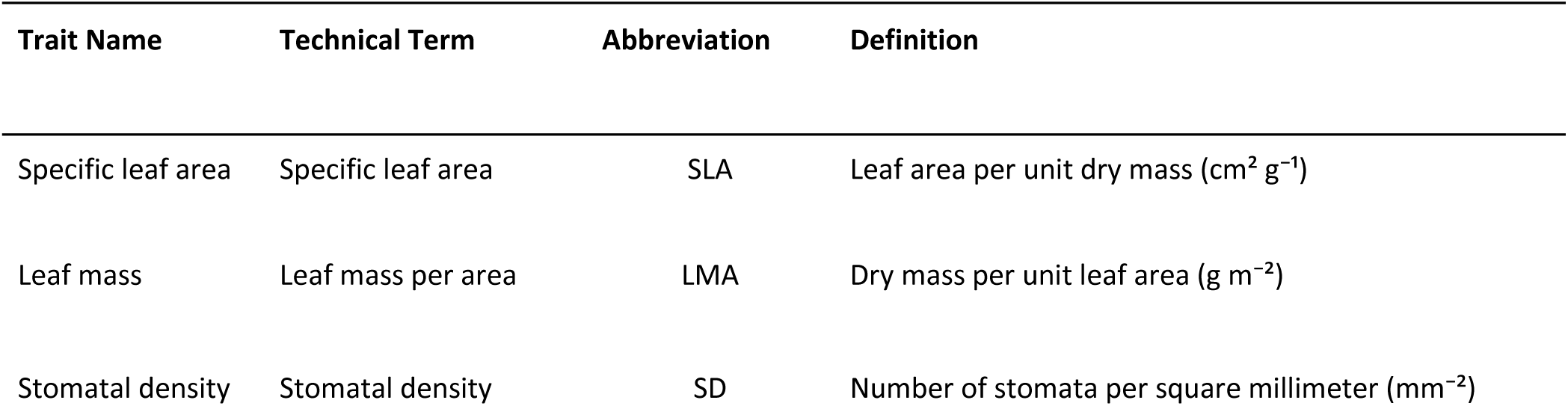
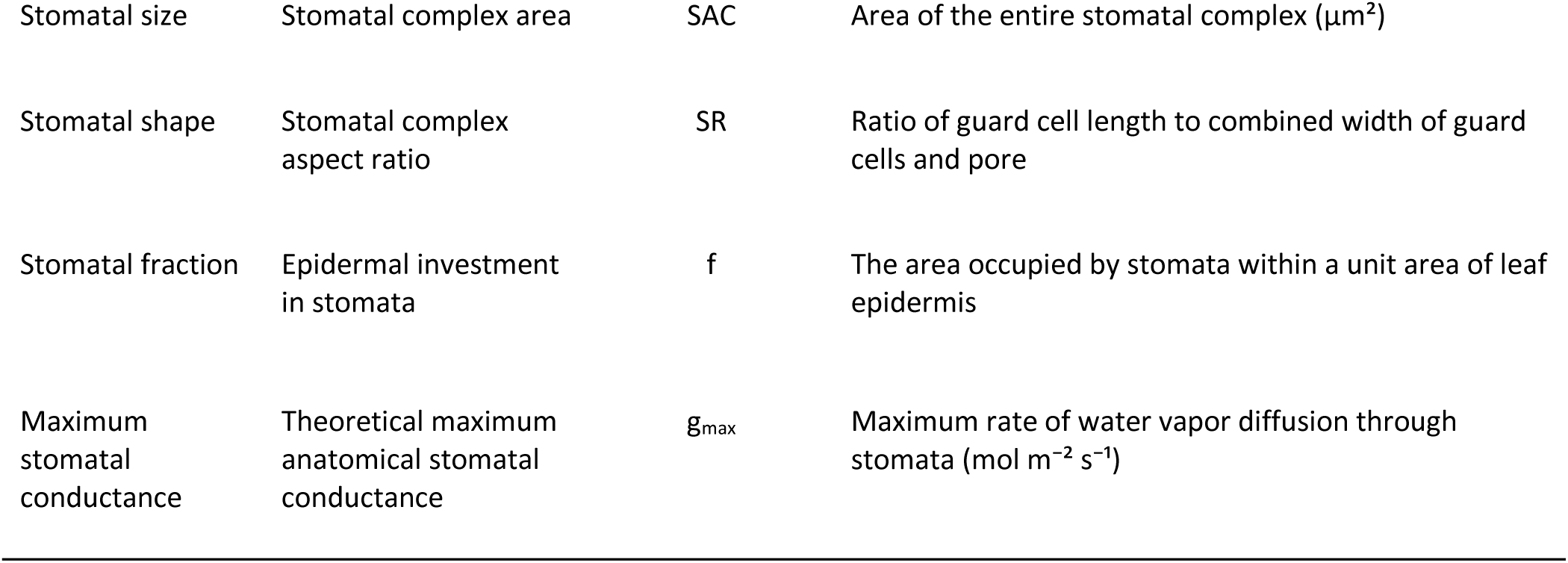
Leaf and stomatal traits measured in this study.

We conducted all analyses in R v4.3.1 (R Core Team, 2025). We first assessed pairwise correlations among all response traits and excluded highly correlated variables (|r| ≥ 0.7; Fig. S3) to avoid redundancy. From each correlated pair, we retained a single representative trait, resulting in a final set comprising SD, SAC, SR, g_max_, SLA and LMA. We did the same for the explanatory variables (Fig. S4). We converted specimen collection dates to day of year (DOY) and calculated photoperiod from DOY and latitude using standard astronomical functions in R (Forsythe et al., 1995), estimating daily daylight hours at each collection site. To summarize the combined influence of multiple, highly correlated climate variables, we performed a principal component analysis (PCA) on the following predictors: mean growing-season temperature (T_gs), maximum temperature (Tmax_gs), potential evapotranspiration (PET_gs) and vapor pressure deficit (VPD_gs). The first principal component (PC1) captured 88% of the total variation, which we used as a composite index of thermal and evaporative conditions. High PC1 values corresponded to warmer climates with high evaporative demand, while low PC1 values indicated cooler climates with lower evaporative demand. We included precipitation (P_gs) as a separate predictor in all models to capture the independent effect of water availability.

We tested for temporal trends, response to CO_2_ and climatic sensitivity in leaf and stomatal traits using generalized additive models (GAMs) implemented in the *mgcv* package (Wood 2017) in R. Prior to the analysis, we standardized all trait variables (z-scored), to enable comparison of effect sizes across traits. Collection year, CO2 and climate variables (PC1_temp and P_gs) were mean-centered prior to the analysis to improve interpretability of interaction terms and reduce collinearity between main effects and interactions. For each trait, we first tested whether responses to collection year, atmospheric CO_2_ or climate were linear or nonlinear. We fitted generalized additive models (GAMs) including a smooth term for each explanatory variable and evaluated the effective degrees of freedom (EDF) to assess response shape. EDF values near 1 indicated approximately linear relationships, whereas higher values indicated increasing curvature in the fitted smooth. Following common conventions (Zuur et al., 2009), we classified relationships as approximately linear when EDF ≤ 1.5, weakly non-linear when EDF was between 1.5 and 2, and strongly non-linear when EDF > 2, provided that the corresponding smooth terms were statistically significant (p < 0.05). We modeled traits showing evidence of linear trends with taxa-specific slopes, whereas we modeled those with non-linear trends with taxa-specific smooth functions of year, CO2 or climate. We included additional smooth terms for spatial coordinates (longitude and latitude) to account for spatial autocorrelation in all models. In addition, we included a smooth term for day-of-year (DOY), to account for phenological variation among specimens for the temporal and CO2 models. We plotted predictions alongside raw standardized trait values and visualized uncertainty using 95% confidence intervals derived from standard errors of predicted values. We assessed model assumptions using diagnostic plots produced with the appraise() function from the *gratia* package (Simpson 2024) in R. We examined residuals versus linear predictor values to evaluate heteroscedasticity and remaining structure, and observed versus fitted values to assess overall model fit. We further inspected a histogram of residuals to check consistency with the assumed error distribution, and a Q–Q plots to evaluate agreement between observed and theoretical residual distributions. We summarized model performance using the percentage of deviance explained as a measure of goodness-of-fit. We initially tested for a habitat-of-origin effect on the measured traits, but as it was not significant, we excluded this predictor from the final models (Fig. S1).

Finally, we used a correlation network approach in R to test for correlation between stomatal and leaf traits. We first z-score standardized all our traits (SLA, LMA, SD, SAC, SR and g_max_), then calculated Pearson correlation coefficients among all trait pairs. We retained only correlations with an absolute value ≥ 0.4. We visualized the resulting network using the Fruchterman–Reingold layout algorithm in the *igraph* package (Csárdi & Nepusz 2006), with edge color indicating the direction of correlation (blue for positive, red for negative) and edge width scaled to correlation strength. Because the relationships among leaf and stomatal traits were consistent between *R. japonica* and *R.* × *bohemica*, we pooled trait data from both taxa to construct a single trait correlation network.

## Results

### Temporal trends

Baseline mean trait values differed between the two taxa. Compared with the hybrid *R.* × *bohemica*, *R. japonica* exhibited lower specific leaf area (F = 29.5, p < 0.001, -16%), higher leaf mass per area (F = 25.7, p < 0.001; +15%), greater stomatal density (F = 4.23, p = 0.03; +5%), larger stomatal complex area (F = 4.04, p = 0.04; +5%) and higher anatomical stomatal conductance (F = 8.69, p = 0.003; +8%) (Fig. S5). Among the six traits measured, specific leaf area, leaf mass per area and stomatal aspect ratio followed predominantly linear trends across years, whereas stomatal density, stomatal complex area and anatomical stomatal conductance showed nonlinear responses for both species (Fig. 2, Table S1). Stomatal complex area changed significantly over time in both taxa, following a nonlinear pattern (EDF > 3, p < 0.05). However, the direction of change differed between taxa: in *R. japonica*, stomatal complex area generally increased over time, whereas in the hybrid *R.* × *bohemica* generally decreased. Stomatal aspect ratio showed a significant taxon-specific temporal slope (F = 4.35, p = 0.037), with a positive trend in *R. japonica* and no significant trend in *R.* × *bohemica*. The remaining traits did not show significant temporal trends in either taxon, although stomatal density in *R. japonica* exhibited a marginally significant negative temporal trend (F = 2.05, p = 0.098).

**Figure 2.**
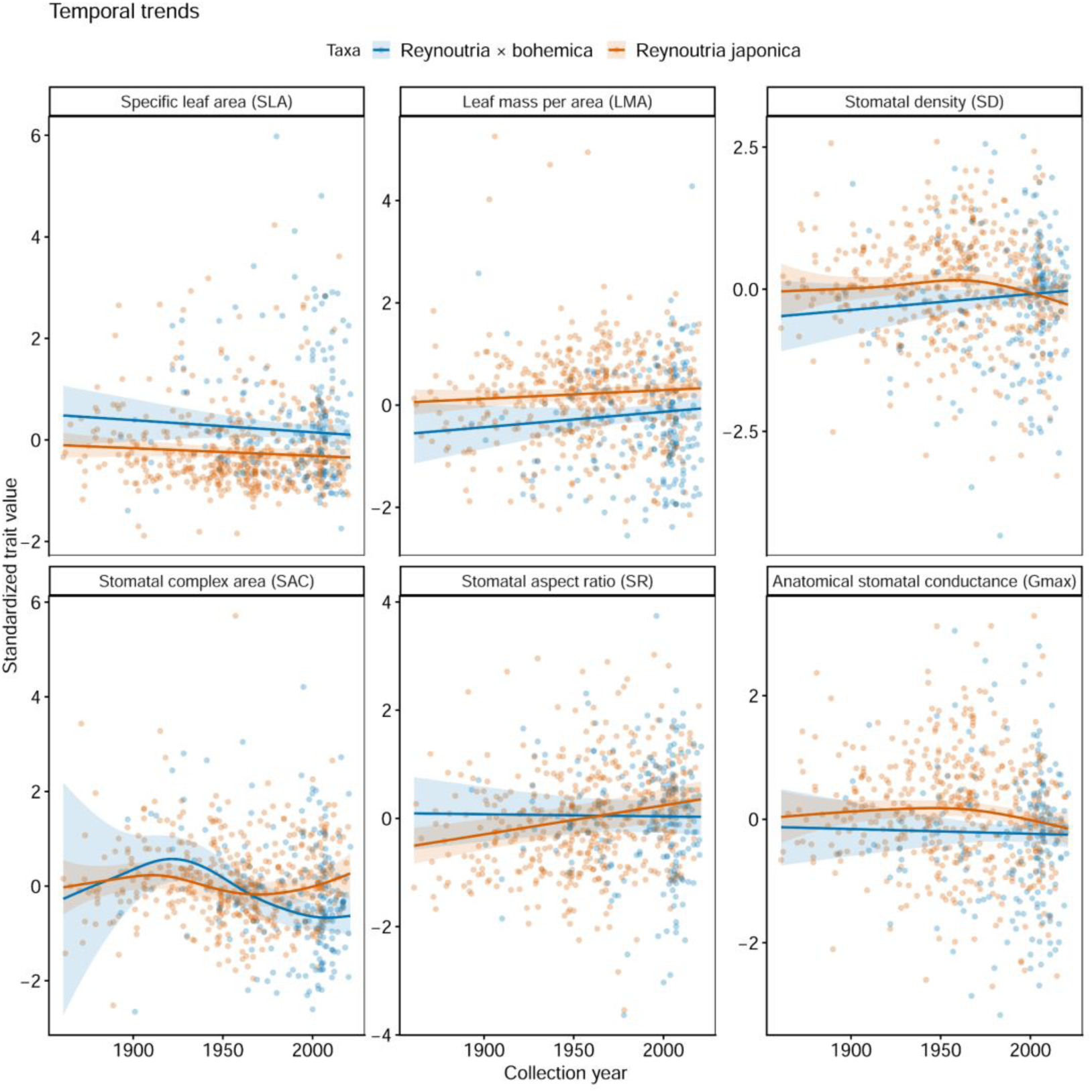
Temporal trends in plant traits. Points represent standardized trait values for individual samples, colored by taxa. Solid lines show GAM-predicted trends over time, with shaded ribbons indicating 95% confidence intervals.

### CO2 effect

Morphological allocation traits (specific leaf area and leaf mass per area), as well as maximum anatomical stomatal conductance, showed weak linear associations with CO_2_ (Fig. 3, Table S2). Stomatal complex area showed a nonlinear association with CO_2_, generally decreasing across the historical CO_2_ gradient in the hybrid *R.* × *bohemica* (F = 7.67, p < 0.001), but showing no significant change in *R. japonica*. *Reynoutria japonica* showed an increase in stomatal ratio associated with elevated CO_2_ (F = 19.41, p < 0.011), whereas *R.* × *bohemica* exhibited no change (Fig. 3). Stomatal density and maximum stomatal conductance showed no significant association with CO_2_, although stomatal density in *R. japonica* showed a modest negative trend (F = 2.53, p = 0.06, Table S2).

**Figure 3.**
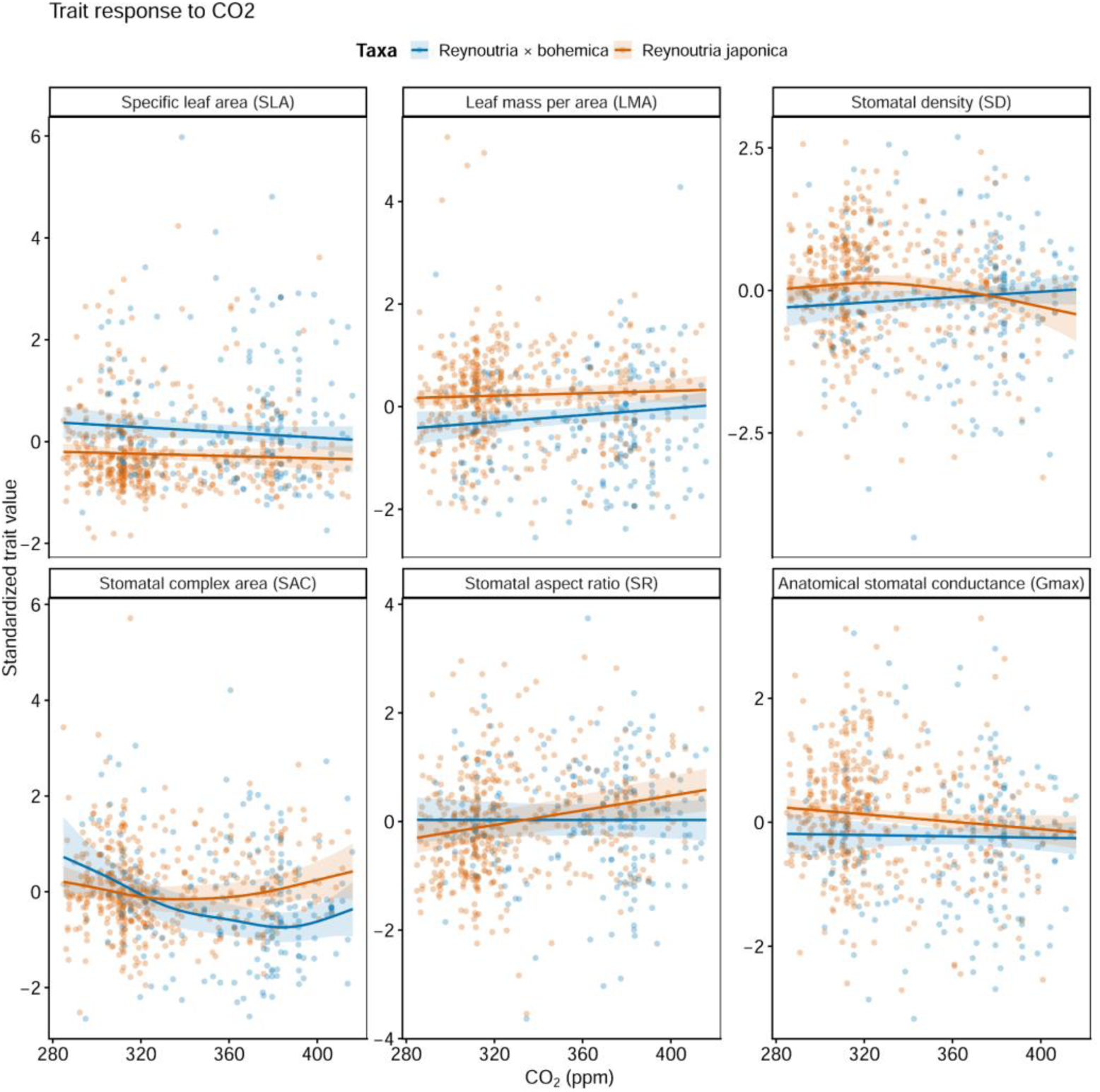
Trait associations with atmospheric CO2. Points represent standardized trait values for individual samples, colored by taxa. Solid lines show GAM-predicted trends over historical CO2, with shaded ribbons indicating 95% confidence intervals.

### Climate sensitivity

Climate effects were strongly trait dependent. We observed nonlinear associations with temperature and evaporative demand (PC1) for specific leaf area, leaf mass per area, stomatal density and stomatal complex area. In contrast, stomatal aspect ratio and maximum anatomical stomatal conductance showed no significant associations with climate variables (Fig. 4, Table S3). Under warmer conditions with high evaporative demand, we found a weak association of reduced SLA in *R. japonica* (EDF = 1.26, p = 0.047), whereas stomatal complex area decreased in both taxa (F = 2.23, p < 0.001). Moreover, stomatal density (F = 1.17, p = 0.0001) and leaf mass per area (EDF = 1.94, p = 0.0001) increased with these conditions in both taxa. In addition, the stomatal area complex (F = 11.1, p < 0.001) increased with increases in the precipitation of the growing season.

**Figure 4.**
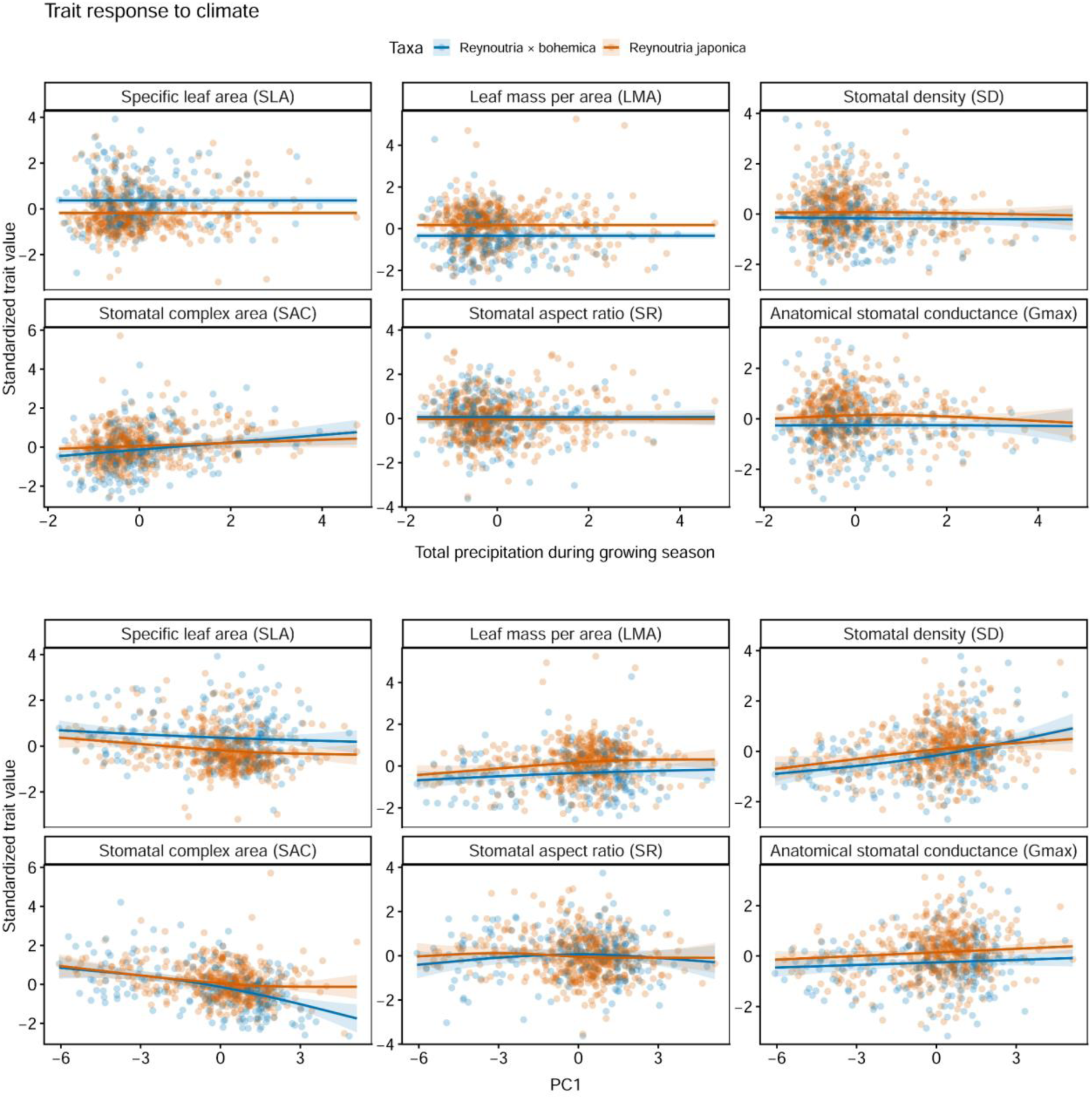
Trait associations with climate variables. Points represent standardized trait values for individual samples, colored by taxa. Solid lines show GAM-predicted trends over precipitation of the growing season and PC1 (heat and evapotranspiration gradient), with shaded ribbons indicating 95% confidence intervals.

### Trait coordination

The trait correlation network revealed a coordinated module linking leaf structural and stomatal traits. Specific leaf area (SLA) was negatively associated with stomatal density (SD) and maximum anatomical stomatal conductance (g_max_), indicating that thinner, resource-acquisitive leaves tend to be associated with reduced stomatal investment and gas exchange capacity. In contrast, leaf mass per area (LMA) showed positive associations with stomatal density and stomatal conductance, reflecting coordinated increases in structural investment and stomatal function. Stomatal complex area (SAC) was negatively correlated with stomatal density, indicating a trade-off between stomatal size and number. Stomatal aspect ratio (SR) showed no strong connections with the trait network (Table S4), suggesting it tends to vary independently of the main leaf–stomatal coordination axis.

**Figure 5.**
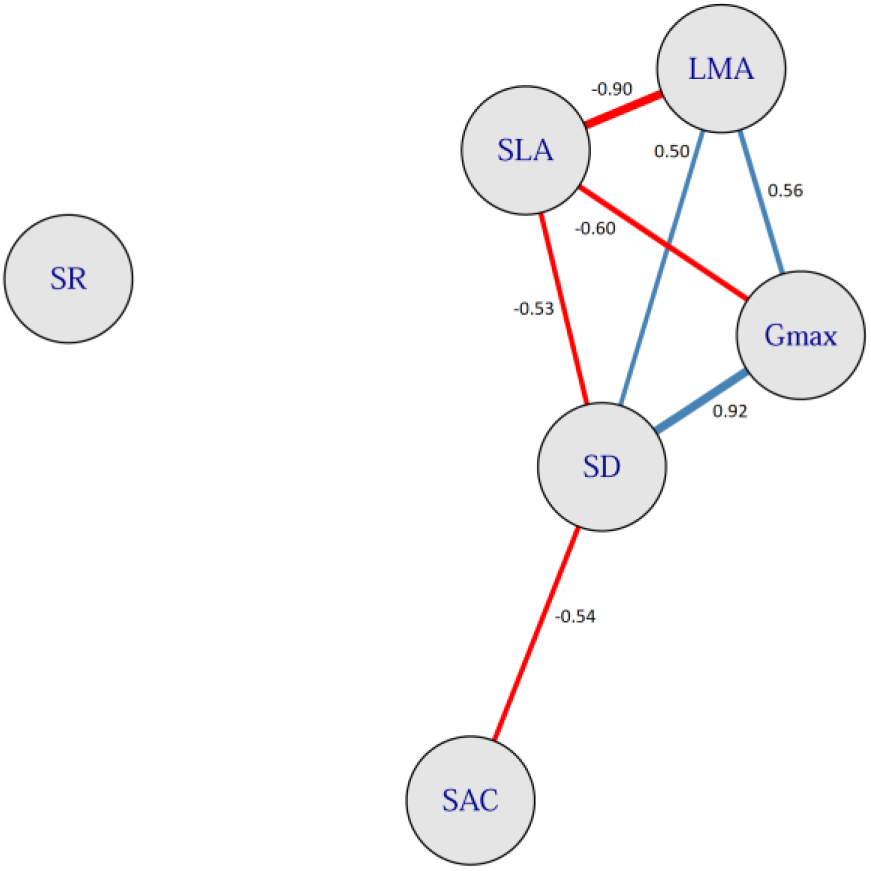
Plant trait correlation network. Nodes represent functional traits and edges represent pairwise correlations between traits. Blue lines indicate positive correlations, and red lines indicate negative correlations, with line thickness proportional to correlation strength. Only statistically significant correlations (Bonferroni-adjusted p < 0.05) above 0.4 are shown.

## Discussion

We used herbarium specimens to examine long-term physiological changes in Japanese knotweed (*R. japonica*) and Bohemian knotweed (*R.* × *bohemica*) across their European invaded range. By linking leaf traits to atmospheric CO_2_ levels as well as to climate variation, we found that both taxa adjusted stomatal traits over time and with rising CO_2_, though their responses differed. Stomatal traits were more strongly associated with temperature and evaporative demand than with precipitation. Leaf structure also constrained function: thin, high-SLA leaves had lower gas-exchange capacity, and stomata exhibited a size–number trade-off. These findings provide an integrated view of how long-term environmental change shapes trait expression in invasive plants and the coordination and trade-offs among functional traits.

We found that leaf traits and most stomatal traits remained stable over time, contrasting with previous studies that reported declines in stomatal density with rising atmospheric CO_2_ across Mediterranean trees, shrubs and herbs (Peñuelas and Matamala, 1990), as well as in large experimental and fossil datasets spanning hundreds of species (Royer, 2001; Large et al., 2017). Local environmental changes often have a stronger effect than global CO2 trends. In addition, plants can respond by adjusting both their stomatal size and number, rather than stomatal density alone (Reid et al., 2003).

Opposite to the direction predicted, *R. japonica* showed an increase in stomatal area complex over time and with increasing CO2, with stomata also becoming more elongated as CO2 increased. Stomatal density showed a weak negative trend with increasing CO2, suggesting some compensation between stomatal number and size (Haworth et al., 2023). In contrast, *R.* × *bohemica* showed a decrease in stomatal area complex over time and with increasing historical CO2 levels. The hybrid also displayed a more acquisitive leaf economic strategy than *R*. *japonica*, characterized by larger, thinner leaves and lower stomatal capacity. This variation in eco-physiological traits within *Reynoutria* could be associated with competitive advantages across different environments. Although both taxa occupy similar habitats in their introduced range—such as riparian zones, roadsides and disturbed areas—the hybrid is generally more robust and capable of exploiting a broader range of environments than its parental species (Jovanović et al., 2018). Smaller stomata in the hybrid may allow rapid adjustment of gas exchange by initiating rapid opening and closing of the stomata which could improve plant resistance to drought and allow closer tracking of environmental variation (Drake et al., 2013). In fact, *Reynoutria* × *bohemica* appears to be expanding into southeastern Europe and the Mediterranean, which may be linked to its relatively higher drought tolerance compared to the parental species (Bailey & Wisskirchen 2004; Jovanović et al., 2018). One limitation of this study is that we could not account for leaf shrinkage, which may have reduced guard cell size and lead to an overestimation of stomatal density. Tests in our lab on fresh and two-week dried *Reynoutria japonica* leaves showed an average leaf shrinkage of approximately 13% (Badreldin, unpublished). However, our measurements (average stomatal length = 35 μm, stomatal width = 25 μm, stomatal density = 100 mm^2^) were consistent with values reported for modern *Reynoutria* individuals in other studies (Khalil et al., 2020; Marschall and Nemoda, 2025), suggesting that shrinkage effects are unlikely to strongly bias relative trends and should be comparable across herbarium specimens. In addition, the hybrid dataset is more sparsely sampled and disproportionately represented by more recent collection dates, therefore these results should be interpreted with caution.

We observed shifts in leaf and physiological traits in both taxa in response to variation in environmental conditions, especially along gradients of temperature and evaporative demand during the growing season. Temperature influences cell division and elevated temperatures can alter stomatal development (Driesen et al., 2020). In hotter and drier weather, plants tended to exhibit lower specific leaf area (SLA) and produced thicker, denser leaves, resulting in higher leaf mass per area (LMA). These structural modifications were accompanied by higher stomatal density while individual stomata became smaller in size. This combination reflects a conservative water-use strategy as smaller stomata can open and close more rapidly. Increased stomatal density may also help sustain gas exchange and photosynthetic activity despite reduced stomatal size, thereby balancing carbon uptake with water conservation (Magaña Ugarte et al., 2020). Our specimens spanned a broad climatic and latitudinal gradient in Europe ranging from coastal areas in Scandinavia to more southern drier locations in northwestern Spain, southern France and northern Italy. Similar shifts in stomatal characteristics and SLA have been documented in shrub species growing in arid environments such as *Protea repens* (Carlson et al., 2016) and *Dodonaea viscosa* subsp. *angustissima* (Hill et al., 2014), suggesting a broadly convergent response across species to aridity (see also the cross-species meta-analysis combining both herbarium and experimental data by Yan et al., 2017). By comparison, both LMA and stomatal complex area increased with higher precipitation during the growing season. Leaves with higher LMA generally have lower mesophyll conductance, which may be compensated for by larger stomata, potentially helping to maintain stomatal conductance and regulate gas exchange and carbon gain when water is abundant. Stomatal traits determine the structural capacity for gas exchange, whereas short-term variation in gas exchange is regulated physiologically through dynamic stomatal opening in response to hydraulic constraints imposed by water availability. Our observations in herbarium specimens align with experimental studies of knotweed conducted in the US, which showed that the species can persist across moisture conditions through coordinated adjustments in leaf structure (LMA), stomatal conductance, transpiration and hydraulic function, allowing it to maintain carbon uptake even under drought stress (Cummins 2013). This plasticity to adjust its physiology to variations in water availability may contribute to knotweed invasion success across different habitats. However, we extracted climate data for the year of collection of each specimen and the climate of the growing season explained only a small proportion of the variation in leaf and stomatal traits (8–20%). These findings suggested that additional unmeasured factors such as soil properties (Adler-Colvin et al., 2025), nutrient availability, light conditions or other microsite characteristics may also play an important role.

Correlations among plant functional traits are widely used to identify the constraints and trade-offs that shape major plant ecological strategies (Ackerly 2004). Trade-offs among traits reflect how plants balance and optimize resource use. Because both leaf economic and physiological traits influence photosynthesis, they are expected to be closely coordinated. In particular, ‘fast’ leaf economic traits (e.g., high SLA) are typically accompanied by ‘fast’ leaf physiological traits (such as small and dense stomata) (Yin et al., 2018). Our results revealed a trade-off in which “fast” leaf economic traits were associated with “slow” physiological traits—characterized by fewer stomata, which also tended to be larger. Similar findings were reported by Pan et al. (2024) for broad leaf tree species in cold-temperate forests. In knotweed, stomata are concentrated on the lower epidermis, which limits water loss while maintaining gas exchange. This trait combination appears to be optimized to support rapid growth in moist habitats—such as those knotweed typically invades—rather than prioritizing strict water conservation. We also found a trade-off between stomatal size and density, consistent with the widespread negative scaling between these traits observed across many other plant species, which has been linked to biophysical constraints on stomatal packing (Liu et al., 2025).

## Conclusions

Our results show that these invasive *Reynoutria* species maintain a stable maximum anatomical gas-exchange capacity over time despite environmentally driven variation in stomatal morphology. Coordinated changes in stomatal density and size appear to preserve this structural capacity for gas exchange across changing environmental conditions. These responses were embedded within a coordinated trait network linking stomatal anatomy with leaf economic traits, highlighting an integrated strategy that balances anatomical plasticity with functional stability. Future studies should use common garden experiments to distinguish heritable variation from within generation phenotypic plasticity in the responses observed here. Expanding trait-based analyses to include belowground traits (rhizomes) and leaf vein architecture as well as soil attributes could provide a more complete understanding of invasive knotweed’s physiological variation and water-use strategy.

## Acknowledgments

This work was supported by the European Union’s Horizon 2020 research and innovation program under the Marie Skłodowska-Curie IF postdoctoral grant agreement No. 101033168 (to REI), and a grant from the German Ministry of Education and Research (BMBF) and the Baden-Württemberg Ministry of Science as part of the Excellence Strategy of the German Federal and State Governments (to REI). We thank Tatiana Tondini, Sulochana Swathi Kannan, Katharina Breyvogel and Rosa Schmitt for their help with data collection in the lab. We thank the curators of the following herbaria for providing us loans of Japanese knotweed herbarium specimens: BG, BREM, BRNU, C, FI, G, GOET, GB, GJO, HEL, JE, KRAM, L, LG, LD, MA, MPU, MSUN, M, NSW, OXF, P, STU, TU, TROM, TRH, TUR, WAG and WU.

## Data availability

All raw data and code generated during this study will be deposited in Zenodo and made publicly available upon acceptance of the manuscript.

## Notes

### Competing Interest Statement

The authors have declared no competing interest.

