## supplemental for "Long-term stomatal and leaf trait dynamics in invasive knotweeds in Europe - insights from 160 years of herbarium records"

### Supplementary information

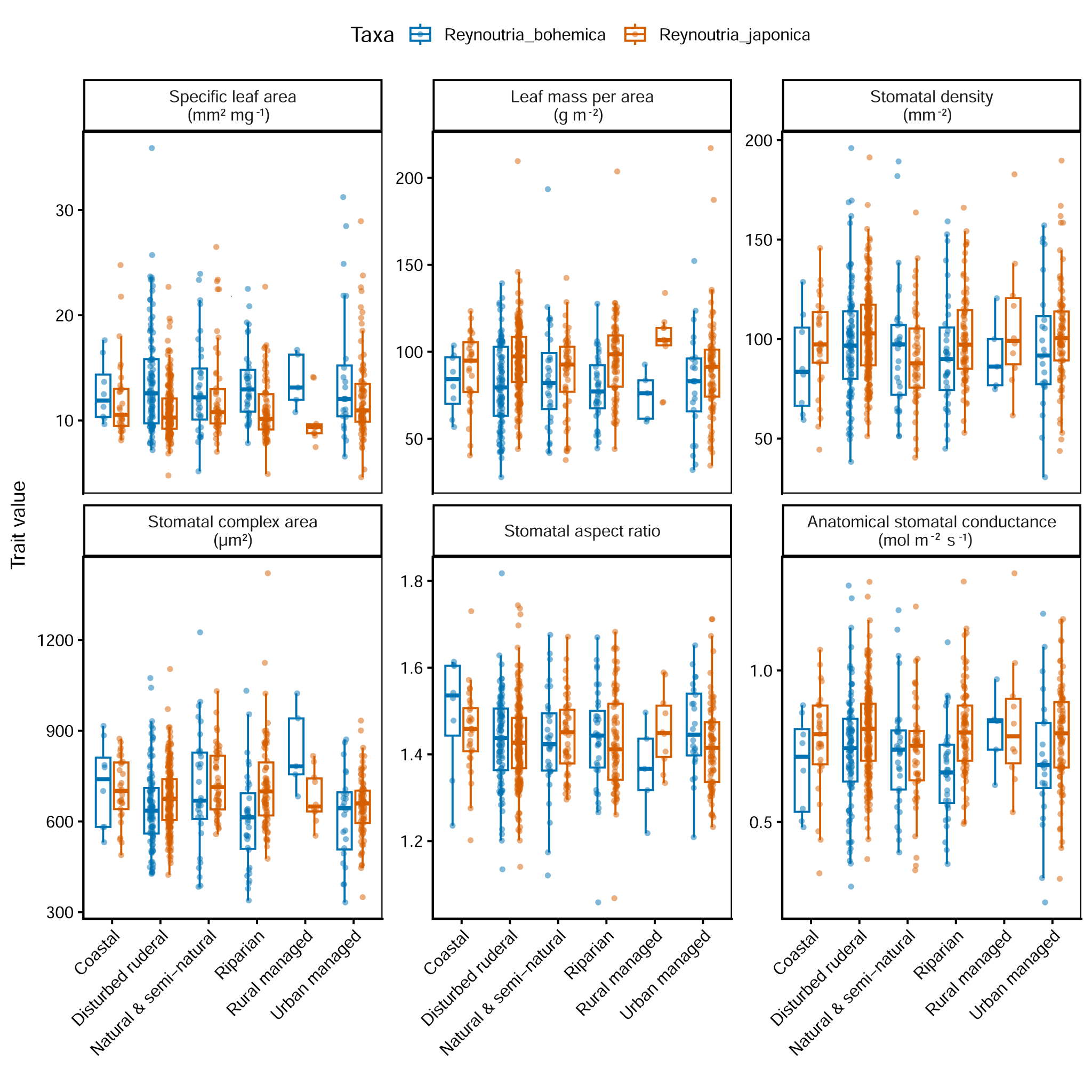
**Figure S1**. Variation in leaf and stomatal traits across major habitat types. Boxplots show the distribution of trait values within each habitat and taxon, with overlaid jittered points representing individual observations.

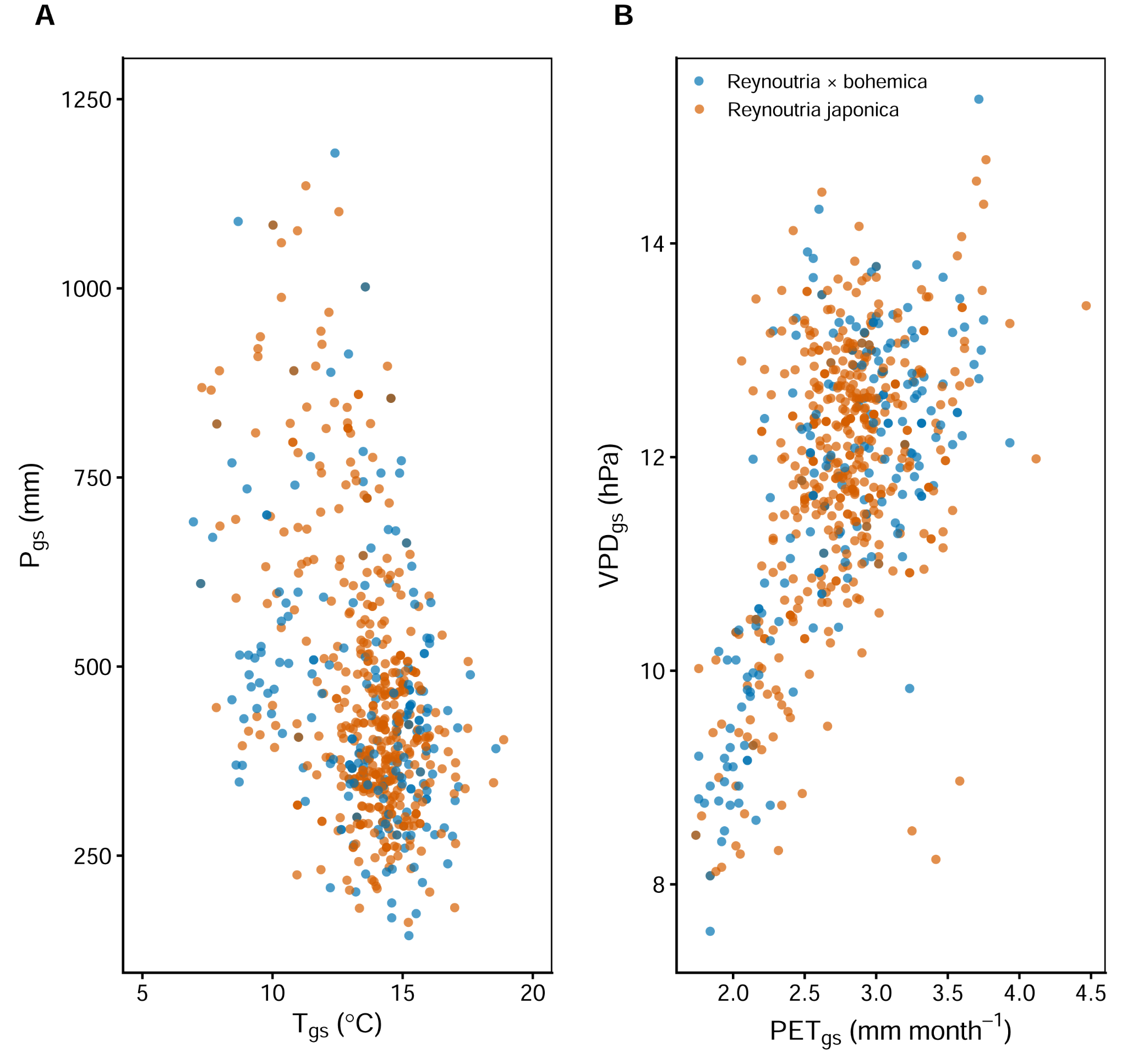

**Figure S2**. Scatterplots showing relationships among climatic variables at each specimen collection site and for each of the two taxa. (A) Total precipitation during the growing season versus average temperature during the growing season and (B) Vapor pressure deficit during growing season versus potential evapotranspiration.

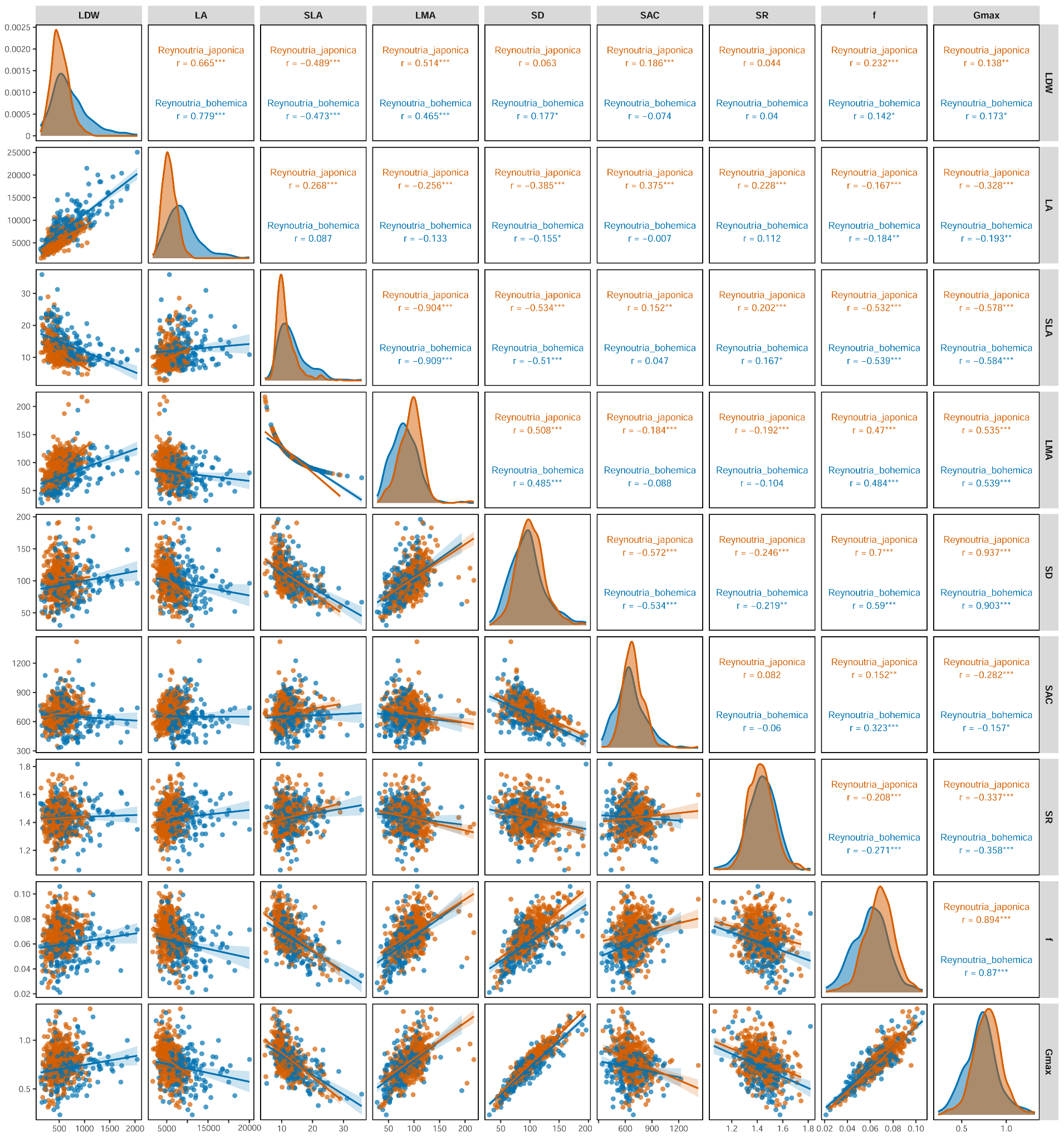

**Figure S3.** Pairwise scatterplot matrix showing correlations among leaf structural and stomatal traits measured in two taxa (*Reynoutria* × *bohemica*: blue; *Reynoutria japonica*: vermilion). Traits include leaf dry weight (LDW), leaf area (LA), specific leaf area (SLA), leaf mass per area (LMA), stomatal density (SD), stomatal complex area (SAC), stomatal ratio (SR), stomatal area fraction (*f*) and theoretical maximum stomatal conductance (Gmax). Diagonal panels show trait distributions (density plots). Lower panels show bivariate scatterplots with fitted linear regression lines and 95% confidence intervals. Upper panels show taxa-specific correlations (Pearson) in corresponding colors. Significance levels are indicated by asterisks (*p* < 0.05, **p* < 0.01, ***p* < 0.001).

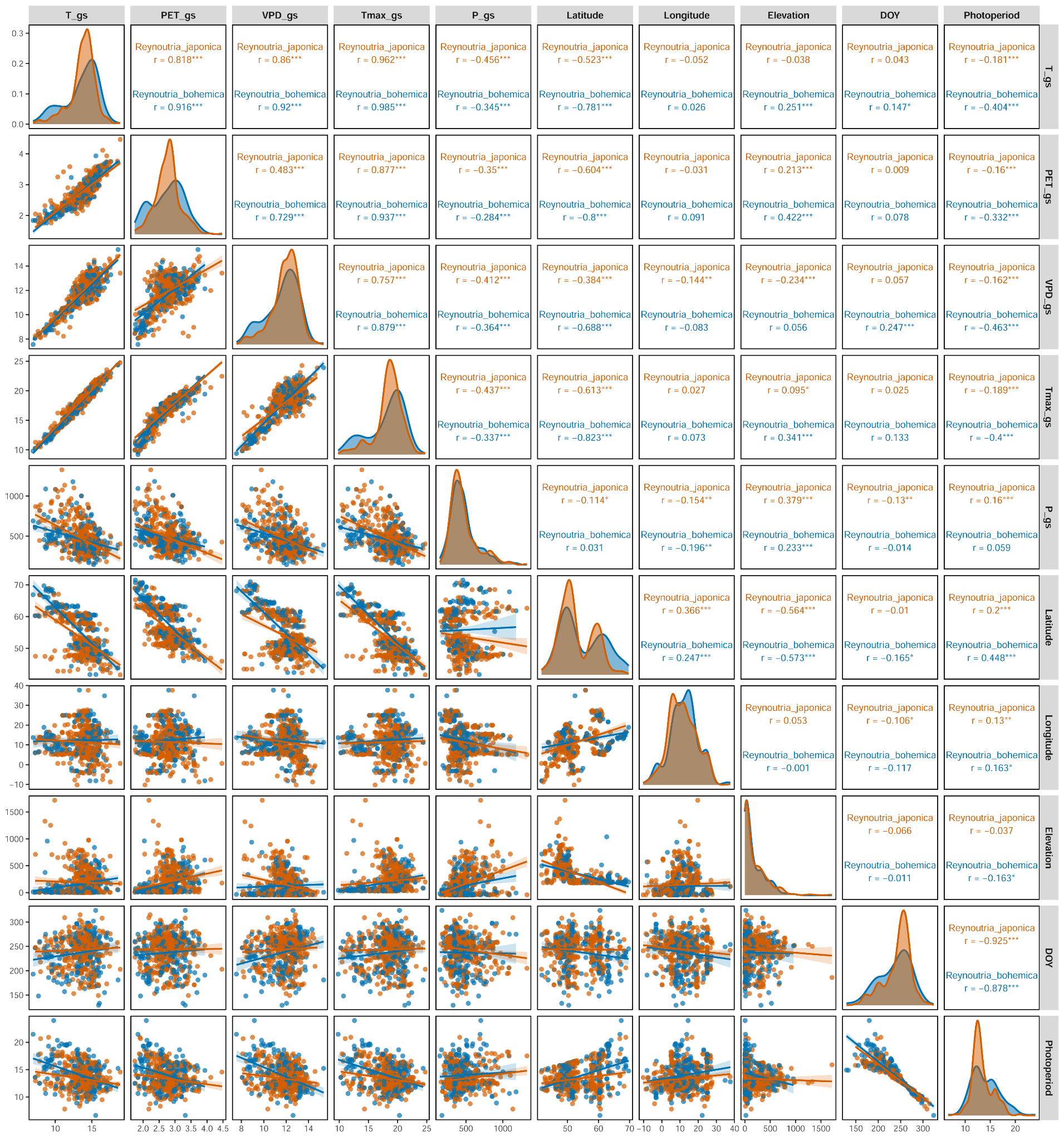

**Figure S4.** Pairwise scatterplot matrix illustrating relationships among environmental variables for herbarium specimen occurrences of *Reynoutria* × *bohemica*: blue; *Reynoutria japonica*: vermilion. Variables include elevation; mean growing-season temperature (T); growing-season potential evapotranspiration (PET); growing-season vapor pressure deficit (VPD); mean maximum growing-season temperature (Tmax); total growing-season precipitation (P), latitude and longitude, elevation, photoperiod (day length) and day of year (DOY). Diagonal panels show kernel density distributions for each variable. Lower panels show bivariate scatterplots with fitted linear regression lines and 95% confidence intervals. Upper panels indicate Pearson correlation coefficients for the two taxa. Asterisks show statistical significance (p < 0.05, *p < 0.01, **p < 0.001).

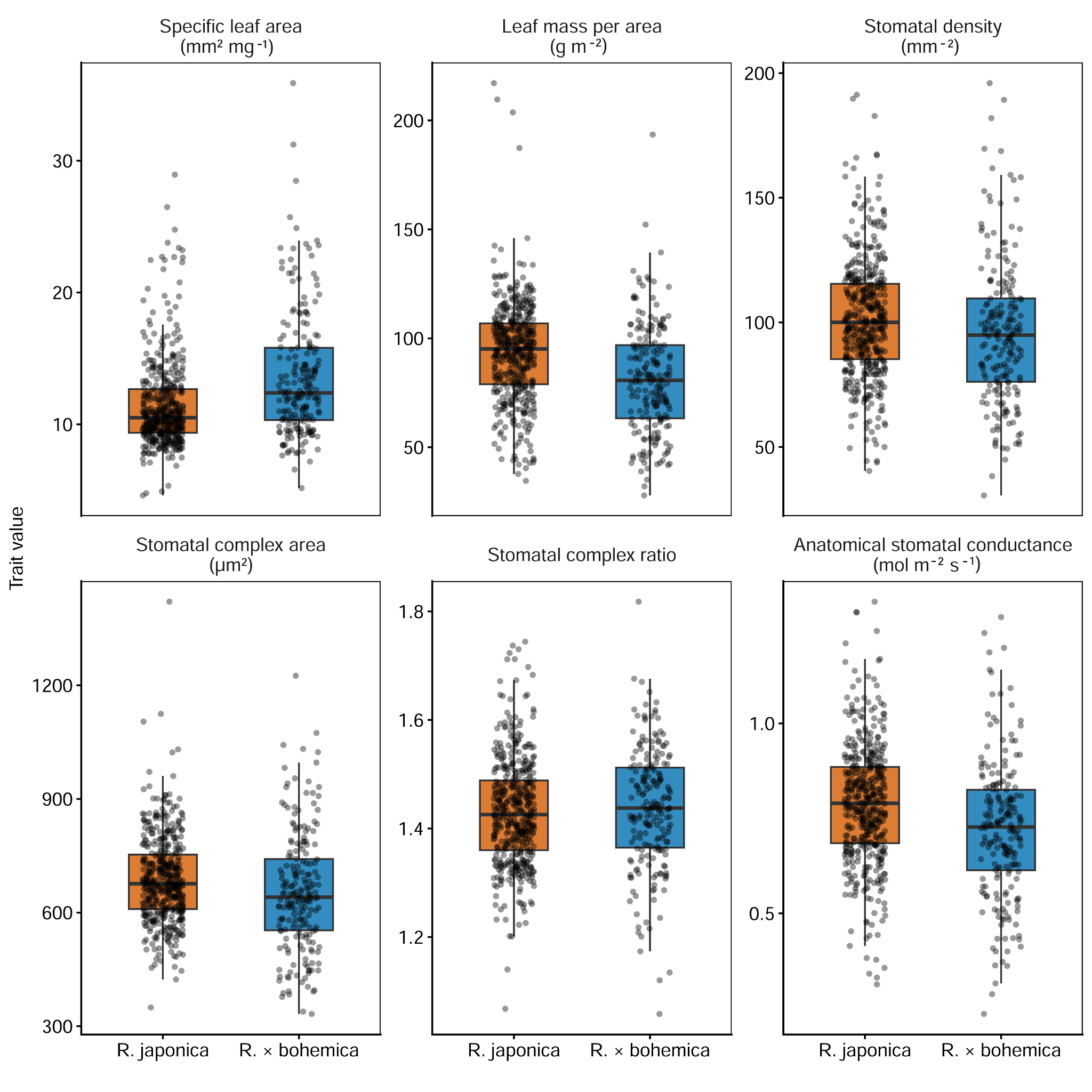

**Figure S5**. Trait variation in *Reynoutria japonica* and *Reynoutria* × *bohemica*. Boxplots show trait distributions for each taxon, with medians, interquartile ranges, whiskers and individual observations (jittered points).

**Table S1.** Summary of generalized additive models (GAMs) for temporal trends in functional traits. The table shows parametric terms (linear effects of taxa, collection year and their interaction) and smooth terms: nonlinear effects of day-of-year s(DOY), spatial coordinates: s(Long, Lat) and species-specific temporal smooths. Abbreviation: df - degrees of freedom for parametric terms, edf - effective degrees of freedom, Ref_df - reference degrees of freedom for smooth terms used in F-tests.

| **Trait** | **Term** | **Type** | **df** | **F-value** | **p-value** | **edf** | **Ref_df** | **Deviance explained** |
| --- | --- | --- | --- | --- | --- | --- | --- | --- |
| SLA | Taxa | parametric | 1 | **29.5** | **< 0.001** | NA | NA | 20.07 |
|  | Collection year | parametric | 1 | 1.08 | 0.298 | NA | NA |  |
|  | Taxa*Collection year | parametric | 1 | 0.11 | 0.730 | NA | NA |  |
|  | s(DOY) | smooth | NA | **27.5** | **< 0.001** | 2.82 | 3.36 |  |
|  | s(Long, Lat) | smooth | NA | **4.65** | **0.009** | 2 | 2 |  |
| LMA | Taxa | parametric | 1 | **25.7** | **< 0.001** | NA | NA | 17.8 |
|  | Collection year | parametric | 1 | 1.71 | 0.191 | NA | NA |  |
|  | Taxa*Collection year | parametric | 1 | 0.26 | 0.605 | NA | NA |  |
|  | s(DOY) | smooth | NA | **23.3** | **< 0.001** | 2.81 | 3.35 |  |
|  | s(Long, Lat) | smooth | NA | **6.17** | **0.002** | 2 | 2 |  |
| SD | s(collection year)*R. × bohemica | smooth | NA | 1.32 | 0.250 | 1 | 1 | 9.04 |
|  | s(collection year)*R. japonica | smooth | NA | 2.05 | 0.098. | 2.6 | 3.25 |  |
|  | s(DOY) | smooth | NA | **11.1** | **< 0.001** | 1 | 1 |  |
|  | s(Long, Lat) | smooth | NA | **10.5** | **< 0.001** | 2.4 | 2.78 |  |
| SAC | s(collection year)*R. × bohemica | smooth | NA | **7.87** | **< 0.001** | 3.22 | 3.98 | 20.9 |
|  | s(collection year)*R. japonica | smooth | NA | **2.65** | **0.027** | 3.61 | 4.46 |  |
|  | s(DOY) | smooth | NA | 0.01 | 0.955 | 1 | 1.01 |  |
|  | s(Long, Lat) | smooth | NA | **6.21** | **< 0.001** | 12 | 16.1 |  |
| SR | Taxa | parametric | 1 | 0.02 | 0.878 | NA | NA | 13.98 |
|  | Collection year | parametric | 1 | 0.02 | 0.873 | NA | NA |  |
|  | Taxa*Collection year | parametric | 1 | **4.34** | **0.037** | NA | NA |  |
|  | s(DOY) | smooth | NA | 1.18 | 0.277 | 1 | 1 |  |
|  | s(Long, Lat) | smooth | NA | **3.06** | **< 0.001** | 15.7 | 20.3 |  |
| g_max_ | s(collection year)*R. × bohemica | smooth | NA | 0.1 | 0.75 | 1 | 1 | 9.09 |
|  | s(collection year)*R. japonica | smooth | NA | 1.94 | 0.16 | 2.09 | 2.63 |  |
|  | s(DOY) | smooth | NA | **12.5** | **< 0.001** | 1 | 1 |  |
|  | s(Long, Lat) | smooth | NA | **8.15** | **< 0.001** | 2 | 2 |  |

**Table S2**. Summary of generalized additive models (GAMs) for taxa responses to CO2. The table shows parametric terms (linear effects of taxa, CO2 and their interaction) and smooth terms: nonlinear effects of day-of-year s(DOY), spatial coordinates: s(Long, Lat) and species-specific CO2 smooths.

| **Trait** | **Term** | **Type** | **df** | **F-value** | **p-value** | **edf** | **Ref_df** | **Deviance explained** |
| --- | --- | --- | --- | --- | --- | --- | --- | --- |
| SLA | Taxa | parametric | 1 | **31.58** | **< 0.001** | NA | NA | 20.3 |
|  | CO2 | parametric | 1 | 1.71 | 0.19 | NA | NA |  |
|  | Taxa*CO2 | parametric | 1 | 0.371 | 0.543 | NA | NA |  |
|  | s(DOY) | smooth | NA | **27.54** | **< 0.001** | 2.82 | 3.36 |  |
|  | s(Long, Lat) | smooth | NA | **4.51** | **0.011** | 2.001 | 2.001 |  |
| LMA | Taxa | parametric | 1 | **27.81** | **< 0.001** | NA | NA | 17.7 |
|  | CO2 | parametric | 1 | 2.814 | 0.093. | NA | NA |  |
|  | Taxa*CO2 | parametric | 1 | 0.775 | 0.378 | NA | NA |  |
|  | s(DOY) | smooth | NA | 23.45 | **< 0.001** | 2.8 | 3.35 |  |
|  | s(Long, Lat) | smooth | NA | 6.082 | **0.002** | 2 | 2 |  |
| SD | s(CO2)*R. × bohemica | smooth | NA | 1.33 | 0.248 | 1 | 1 | 8.91 |
|  | s(CO2)*R. japonica | smooth | NA | 2.53 | 0.069. | 2.1 | 2.65 |  |
|  | s(DOY) | smooth | NA | 11.57 | **< 0.001** | 1 | 1 |  |
|  | s(Long, Lat) | smooth | NA | 11.99 | **< 0.001** | 2.21 | 2.41 |  |
| SAC | s(CO2)*R. × bohemica | smooth | NA | **7.67** | **< 0.001** | 3.03 | 3.76 | 20.4 |
|  | s(CO2)*R. japonica | smooth | NA | **2.29** | **0.07.** | 2.65 | 3.31 |  |
|  | s(DOY) | smooth | NA | 0.31 | 0.68 | 2.16 | 2.68 |  |
|  | s(Long, Lat) | smooth | NA | **6.24** | **< 0.001** | 12.1 | 16.1 |  |
| SR | s(CO2)*R. × bohemica | smooth | NA | 0 | 0.99 | 1 | 1 | 14.1 |
|  | s(CO2)*R. japonica | smooth | NA | **19.41** | **< 0.001** | 1 | 1 |  |
|  | s(DOY) | smooth | NA | 0.98 | 0.32 | 1 | 1 |  |
|  | s(Long, Lat) | smooth | NA | **3.13** | **< 0.001** | 15.38 | 20.02 |  |
| g_max_ | Taxa | parametric | 1 | **9.43** | **0.002** | NA | NA | 8.86 |
|  | CO2 | parametric | 1 | 0.06 | 0.79 | NA | NA |  |
|  | Taxa*CO2 | parametric | 1 | 0.95 | 0.32 | NA | NA |  |
|  | s(DOY) | smooth | NA | 12.87 | **< 0.001** | 1 | 1 |  |
|  | s(Long, Lat) | smooth | NA | 7.85 | **< 0.001** | 2.002 | 2.005 |  |

**Table S3.** Summary of generalized additive models (GAMs) for taxa responses to climate. The table shows parametric terms (linear effects of taxa and precipitation) and smooth terms: nonlinear effects of s(Long, Lat) and taxa-specific PC1 smooths.

| Trait | Term | Type | df | F-value | p-value | edf | Ref_df | Deviance explained |
| --- | --- | --- | --- | --- | --- | --- | --- | --- |
| SLA | Taxa | parametric | 1 | **42.67** | **< 0.001** | NA | NA | 8.9 |
|  | P_gs | parametric | 1 | 0.268 | 0.605 | NA | NA |  |
|  | s(PC1)*R. × bohemica | smooth | NA | 0 | 0.508 | 0.0006 | 9 |  |
|  | s(PC1)*R. japonica | smooth | NA | **0.393** | **0.047** | 1.269 | 9 |  |
|  | s(Long, Lat) | smooth | NA | **0.14** | **0.043** | 1.323 | 29 |  |
| LMA | Taxa | parametric | 1 | **36.45** | **< 0.001** | NA | NA | 8.38 |
|  | P_gs | parametric | 1 | 0.201 | 0.654 | NA | NA |  |
|  | s(PC1)*R. × bohemica | smooth | NA | **0.325** | **0.047** | 0.746 | 9 |  |
|  | s(PC1)*R. japonica | smooth | NA | **1.46** | **< 0.001** | 1.937 | 9 |  |
|  | s(Long, Lat) | smooth | NA | 0 | 0.53 | 0.00007 | 29 |  |
| SD | Taxa | parametric | 1 | **4.26** | **0.039** | NA | NA | 9.01 |
|  | P_gs | parametric | 1 | **5.81** | **0.016** | NA | NA |  |
|  | s(PC1)*R. × bohemica | smooth | NA | **1.19** | **< 0.001** | 1.74 | 9 |  |
|  | s(PC1)*R. japonica | smooth | NA | **0.31** | **0.03** | 0.746 | 9 |  |
|  | s(Long, Lat) | smooth | NA | **0.182** | **0.022** | 1.496 | 29 |  |
| SAC | Taxa | parametric | 1 | **12.6** | **< 0.001** | NA | NA | 18.1 |
|  | P_gs | parametric | 1 | **11.09** | **< 0.001** | NA | NA |  |
|  | s(PC1)*R. × bohemica | smooth | NA | **5.26** | **< 0.001** | 2.03 | 9 |  |
|  | s(PC1)*R. japonica | smooth | NA | **1.08** | **0.001** | 1.86 | 9 |  |
|  | s(Long, Lat) | smooth | NA | 0.09 | 0.08 | 1.073 | 29 |  |
| SR | Taxa | parametric | 1 | 0.012 | 0.912 | NA | NA | 10.1 |
|  | PC1 | parametric | 1 | 1.86 | 0.172 | NA | NA |  |
|  | P_gs | parametric | 1 | 0.229 | 0.632 | NA | NA |  |
|  | PC1*Taxa | parametric | 1 | 3.467 | 0.063 | NA | NA |  |
|  | P_gs*Taxa | parametric |  | 0.107 | 0.743 | NA | NA |  |
|  | s(Long, Lat) | smooth | NA | 1.66 | **< 0.001** | 11.18 | 29 |  |
| g_max_ | Taxa | parametric | 1 | 19.28 | **< 0.001** | NA | NA | 7.19 |
|  | PC1 | parametric | 1 | 1.053 | 0.305 | NA | NA |  |
|  | P_gs | parametric | 1 | 0.295 | 0.587 | NA | NA |  |
|  | PC1*Taxa | parametric | 1 | 0.077 | 0.781 | NA | NA |  |
|  | P_gs*Taxa | parametric | 1 | 0.004 | 0.952 | NA | NA |  |
|  | s(Long, Lat) | smooth | NA | **0.268** | **0.006** | 1.89 | 29 |  |

**Table S4**. Pearson correlation coefficients among leaf (specific leaf area: SLA, leaf mass per area: LMA) and stomatal traits (stomatal density: SD, stomatal area complex: SAC, stomatal aspect ratio: SR and maximum anatomical stomatal conductance: g_max_). Only statistically significant correlations (Bonferroni-adjusted p < 0.05) are shown. Positive values indicate positive relationships between traits, while negative values indicate trade-offs.

| Trait | SLA | LMA | SD | SAC | SR | g_max_ |
| --- | --- | --- | --- | --- | --- | --- |
| SLA | 1 | -0.90 | -0.52 | - | 0.18 | -0.59 |
| LMA | -0.90 | 1 | 0.50 | - | -0.16 | 0.55 |
| SD | -0.52 | 0.50 | 1 | -0.53 | -0.23 | 0.92 |
| SAC | - | - | -0.53 | 1 | - | -0.20 |
| SR | 0.18 | -0.16 | -0.23 | - | 1 | -0.34 |
| g_max_ | -0.59 | 0.55 | 0.92 | -0.20 | -0.34 | 1 |
